# Extraction method dominates phytochemical profiles of aromatic medicinal herbs: a comparative HPTLC study

**DOI:** 10.64898/2025.12.18.694438

**Authors:** Daniel Faizy, Arvin Azodi, Ahmet Naif Kaya, Farbod Goshtasebi Poor

**Affiliations:** Biomedical Engineering Department, Acibadem University; Department of Chemistry, University of Toronto; Faculty of Medicine, Hacettepe University

## Abstract

The extraction method is a critical determinant of the phytochemical profile obtained from medicinal plants, yet comparative evaluations across multiple extraction techniques and plant species remain limited. In this study, the qualitative influence of commonly used extraction methods on the phytochemical composition of aromatic medicinal herbs was investigated using high-performance thin-layer chromatography (HPTLC). Ten aromatic medicinal herbs belonging to the families *Lamiaceae*, *Apiaceae*, and *Asteraceae* were subjected to six extraction approaches: simple distillation, essential oil (essence) extraction, hydroalcoholic tincture, infusion, decoction, and decoction of distillate. All extracts were analyzed under standardized HPTLC conditions using silica gel 60 F254 plates, a toluene-ethyl acetate (93:7 v/v) mobile phase, anisaldehyde-sulfuric acid derivatization, and a standard as a reference compound. Across all investigated species, the extraction method exerted a stronger influence on band number, intensity, and polarity distribution than botanical differences among the herbs. Essential oil and tincture extracts consistently produced the highest diversity of bands, particularly in higher Rf regions corresponding to non-polar and semi-polar compounds. Aqueous methods, including infusion and decoction, predominantly yielded low-Rf polar bands with reduced overall diversity, while simple distillation showed minimal extraction efficiency. These findings demonstrate that extraction strategy is the primary factor governing qualitative phytochemical outcomes in HPTLC-based herbal analysis. The results highlight the importance of method selection in phytochemical screening and provide a comparative framework for choosing appropriate extraction techniques in pharmacological, cosmetic, and analytical applications.

## 1. Introduction

Medicinal and aromatic plants have long been used in traditional and modern healthcare systems as sources of bioactive compounds with antimicrobial, antioxidant, anti-inflammatory, and therapeutic properties. These bioactivities arise from complex mixtures of secondary metabolites, including terpenoids, phenolics, aldehydes, and alcohols, whose chemical characteristics strongly influence their extraction, stability, and detectability (Hajhashemi et al., 2002; Meeran et al., 2017). Consequently, the selection of an appropriate extraction method represents a critical step in phytochemical investigations, as it governs both the qualitative and quantitative composition of the resulting extract.

### 1.1 Influence of Extraction Method on Phytochemical Profiles

Extraction techniques differ substantially in solvent polarity, temperature, duration, and mechanical processing, all of which affect the solubility and preservation of plant metabolites. Volatile and lipophilic compounds, such as monoterpenes and phenolic terpenoids (e.g., thymol, menthol, and anethole), are preferentially recovered through non-polar or semi-polar extraction strategies, including essential oil isolation and hydroalcoholic tinctures (Borugă et al., 2014; Basti et al., 2016). In contrast, aqueous extractions such as infusion and decoction predominantly extract polar constituents, including glycosides and water-soluble phenolics, often at the expense of volatile compounds that may be lost through evaporation or thermal degradation (Sharma & Anand, 1997). Despite the widespread use of multiple extraction techniques in herbal medicine and industry, comparative studies that evaluate several extraction methods under standardized analytical conditions remain limited. Many investigations focus on a single extraction approach or a single plant species, making it difficult to generalize conclusions regarding extraction efficiency or phytochemical diversity. As a result, there is a need for systematic comparisons that isolate the effect of extraction method from botanical variability.

### 1.2 Aromatic Medicinal Herbs as a Comparative Model

Aromatic medicinal herbs provide a suitable model system for evaluating extraction methods due to their well-characterized volatile and semi-volatile phytochemical composition. Species belonging to families such as *Lamiaceae*, *Apiaceae*, and *Asteraceae* are particularly rich in essential oils and phenolic terpenoids and are extensively used in traditional medicine, food preservation, cosmetics, and pharmaceutical formulations (Şahin et al., 2003; Kuete, 2017). While botanical differences influence the relative abundance of individual compounds, these herbs share broadly similar classes of metabolites, allowing extraction-dependent effects to be examined across multiple species without confounding chemical incompatibility. By incorporating multiple aromatic herbs from different botanical families, comparative analyses can assess whether extraction method exerts a more pronounced influence on observed phytochemical profiles than species-specific variation. Such an approach strengthens methodological conclusions and enhances the relevance of the findings for applied fields, including herbal drug development and quality control.

### 1.3 Role of HPTLC in Comparative Phytochemical Analysis

High-performance thin-layer chromatography (HPTLC) is a well-established analytical technique for qualitative phytochemical screening and comparative analysis of complex plant extracts. Compared to conventional thin-layer chromatography (TLC), HPTLC offers improved resolution, higher reproducibility, automated sample application, and enhanced visualization, making it particularly suitable for multi-sample comparisons under standardized conditions (Srivastava, 2011; Attimarad et al., 2011). HPTLC enables simultaneous analysis of numerous extracts on a single plate, facilitating direct visual comparison of band number, intensity, and migration behavior (Rf values). When coupled with appropriate derivatization reagents and reference standards, HPTLC provides valuable insight into the polarity distribution and relative diversity of extracted compounds, even in the absence of full compound identification. This makes HPTLC especially appropriate for extraction-method-focused studies where qualitative trends are of primary interest.

### 1.4 Study Objective

The aim of this study was to systematically compare the qualitative effects of commonly used extraction methods on the phytochemical profiles of aromatic medicinal herbs using HPTLC. Multiple plant species from different botanical families were subjected to six extraction techniques: simple distillation, essential oil extraction, hydroalcoholic tincture, infusion, decoction, and decoction of distillate; followed by analysis under standardized HPTLC conditions. By emphasizing extraction method rather than botanical specificity, this study seeks to clarify how methodological choices influence observed phytochemical diversity and polarity, thereby providing a comparative framework for selecting appropriate extraction strategies in analytical, pharmaceutical, and industrial applications.

## 2. Materials and Methods

### 2.1 Plant Materials

Ten aromatic medicinal herbs were investigated in this study, representing three botanical families: Lamiaceae (Mentha longifolia, Mentha piperita, Mentha spicata, Thymus vulgaris, Satureja hortensis, Zataria multiflora), Apiaceae (Foeniculum vulgare, Anethum graveolens), and Asteraceae (Artemisia dracunculus). Plant materials were obtained from local herbal suppliers and botanical sources and were authenticated based on macroscopic characteristics. Leaves or aerial parts were air-dried at ambient temperature in a shaded environment to minimize degradation of volatile compounds. Dried materials were stored in sealed containers prior to extraction.

In addition to the named species, an additional unidentified mint specimen (hereafter “unknown mint”) was collected from Shahr-e Rey and processed identically to the other mint samples for comparative HPTLC analysis.

### 2.2 Extraction Procedures

All herbs were processed independently using six commonly employed extraction methods. To ensure comparability, extraction conditions were standardized across species wherever applicable.

Extraction parameters (plant mass, solvent volume, and extraction duration) were kept consistent within each extraction method across all samples but are reported qualitatively, as the aim of this study was comparative pattern analysis rather than quantitative optimization.

#### 2.2.1 Simple Distillation

Approximately 25 g of dried plant material was immersed in distilled water and subjected to simple distillation using a Clevenger-type apparatus. The mixture was heated at approximately 80 °C for 2 h. The aqueous distillate was collected for analysis, while the residual decoction remaining in the distillation flask was retained as a separate sample.

#### 2.2.2 Essential Oil (Essence) Extraction

Essential oils were collected during the simple distillation process and separated from the aqueous phase based on density differences. The oil fraction was carefully isolated and diluted with hexane prior to chromatographic analysis to ensure appropriate loading volume and band resolution.

#### 2.2.3 Decoction of Distillate

The residual aqueous phase remaining after simple distillation was further concentrated by gentle heating until a viscous extract was obtained. This decoction of distillate was collected, cooled, and stored in sealed containers for subsequent analysis.

#### 2.2.4 Infusion

For infusion, 12.5 g of dried plant material was immersed in 250 mL of boiling distilled water. The mixture was covered to minimize volatilization and allowed to steep for 1 h. Solid plant material was removed by filtration, and the resulting liquid was concentrated by evaporation until a viscous extract was obtained.

#### 2.2.5 Decoction

Decoction was performed by boiling 12.5 g of dried plant material in 250 mL of distilled water for 15 min. After filtration, the extract was concentrated by continued heating to yield a viscous residue suitable for chromatographic analysis.

#### 2.2.6 Tincture

Hydroalcoholic tinctures were prepared by immersing 12.5 g of dried plant material in a 50:50 (v/v) ethanol-water solution. The mixture was sealed and stored in darkness at room temperature for 48 h with intermittent agitation. After filtration, the extract was concentrated by evaporation to obtain a viscous tincture.

### 2.3 Sample Preparation for HPTLC Analysis

Viscous extracts obtained from infusion, decoction, tincture, and decoction of distillate were weighed (0.1 g) and dissolved in 1 mL of methanol. Samples were vortex-mixed, sonicated for 5 min to ensure complete dissolution, and centrifuged at 10,000 rpm for 1 min. Supernatants were transferred to chromatography vials for analysis. Reference standard solutions of thymol, menthol, and anethole were prepared individually by dissolving 0.1 g of each compound in 1 mL of methanol. Each standard was applied to HPTLC plates alongside corresponding plant extracts to facilitate qualitative comparison of migration behavior and band visualization. Essential oil samples were diluted with hexane prior to application to prevent overloading and band distortion.

### 2.4 HPTLC Conditions

HPTLC analysis was performed on silica gel 60 F254 plates (10 × 10 cm). Samples were applied using a microliter pipette, with standardized loading volumes across all extracts. Chromatographic development was carried out in a twin-trough chamber saturated with a mobile phase consisting of toluene and ethyl acetate (93:7 v/v). The development chamber was pre-saturated with mobile-phase vapor for approximately 20 min prior to plate development. After development, plates were air-dried and derivatized using anisaldehyde-sulfuric acid reagent, followed by brief heating to enhance band visualization. Plates were examined under UV and white light, and chromatograms were documented using an HPTLC visualizer.

### 2.5 Data Interpretation

Chromatographic profiles were evaluated qualitatively based on the number of resolved bands, their relative intensity, and migration behavior (Rf values). Particular attention was given to the polarity distribution of compounds, inferred from band position relative to the reference standard. No quantitative densitometric analysis was performed.

### 2.6 Safety and Ethical Considerations

All laboratory procedures were conducted in accordance with standard chemical safety protocols. Solvents and reagents were handled using appropriate personal protective equipment. No human or animal subjects were involved in this study.

## 3. Results

### 3.1 Overall Influence of Extraction Method

HPTLC analysis revealed that the extraction method exerted a dominant influence on the qualitative phytochemical profiles of all investigated herbs. Across species and botanical families, extracts clustered more clearly by extraction technique than by plant identity. Differences among herbs were observed primarily in band intensity and relative prominence rather than in the overall pattern of compound distribution. This indicates that extraction conditions governed the diversity, polarity range, and detectability of compounds to a greater extent than botanical variation.

Comparable extraction-dependent trends were also observed within species: Mentha longifolia samples collected from two geographic locations (Bagh-e Firoozeh and Shahr-e Rey) produced qualitatively similar HPTLC profiles when prepared by the same extraction methods (see Figure 3).

**Figure 1.**
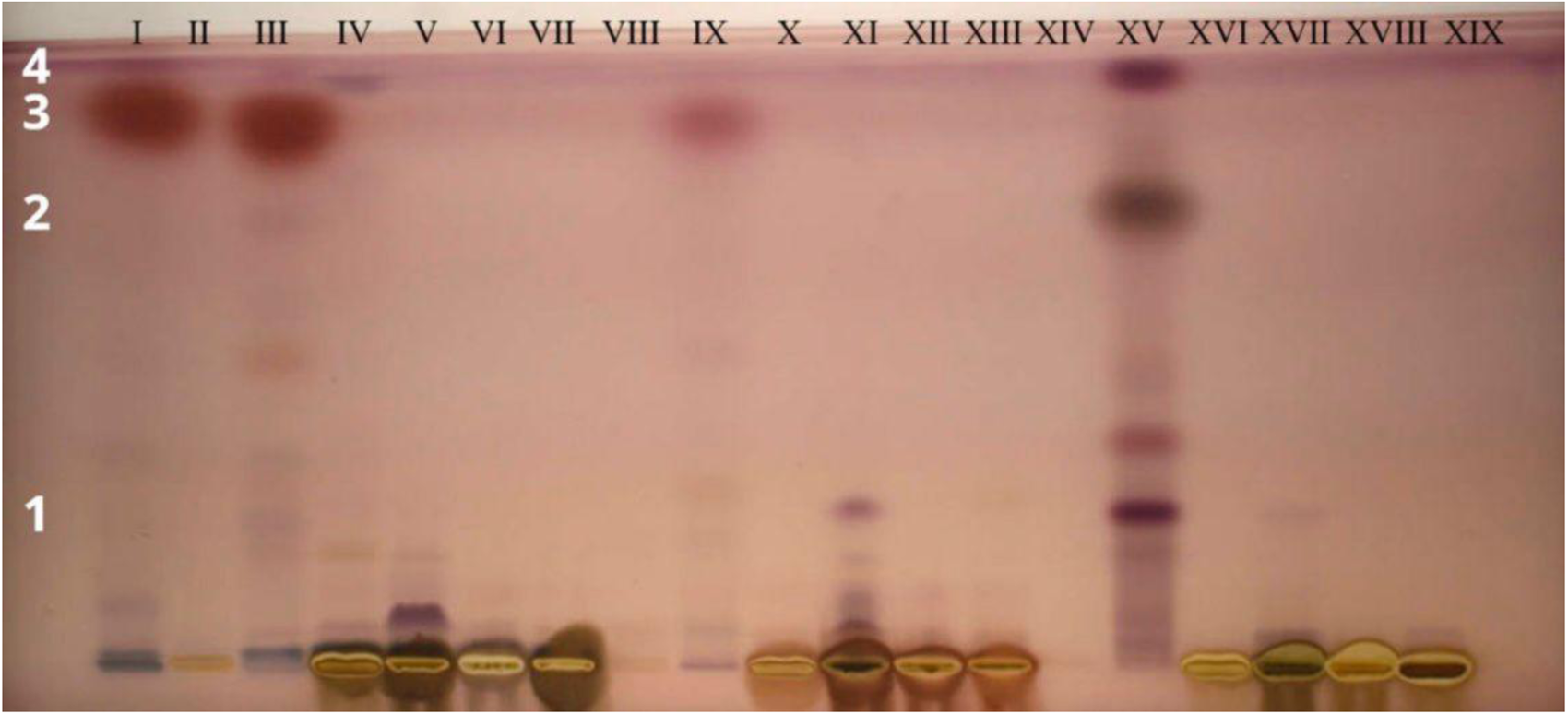
HPTLC analysis of samples. Arabic numerals represent bands, while roman numerals represent samples. I) Standard (Anethole). Foeniculum Vulgare is from Band Column II to VII: II) Simple Distillation III) Essence extracted from Simple Distillation IV) Decoction from Simple Distillation V) Tincture VI) Infusion VII) Decoction. Artemisia Dracunculus is from Band Column VIII to XIII: VIII) Simple Distillation IX) Essence Extracted from Simple Distillation X) Decoction from Simple Distillation XI) Tincture XII) Infusion XIII) Decoction. Anethum Graveolens L. is from Band Column XIV to XIX: XIV) Simple Distillation XV) Essence Extracted from Simple Distillation XVI) Decoction from Simple Distillation XVII) Tincture XVIII) Infusion XIX) Decoction.

An additional sample of unidentified mint from Shahr-e Rey produced HPTLC profiles that closely resembled those of Mentha spicata (see Figure 2, lanes XIV–XIX for unknown mint), suggesting a likely taxonomic affinity with spearmint; however, definitive species identification would require molecular methods such as PCR or DNA barcoding.

**Figure 2.**
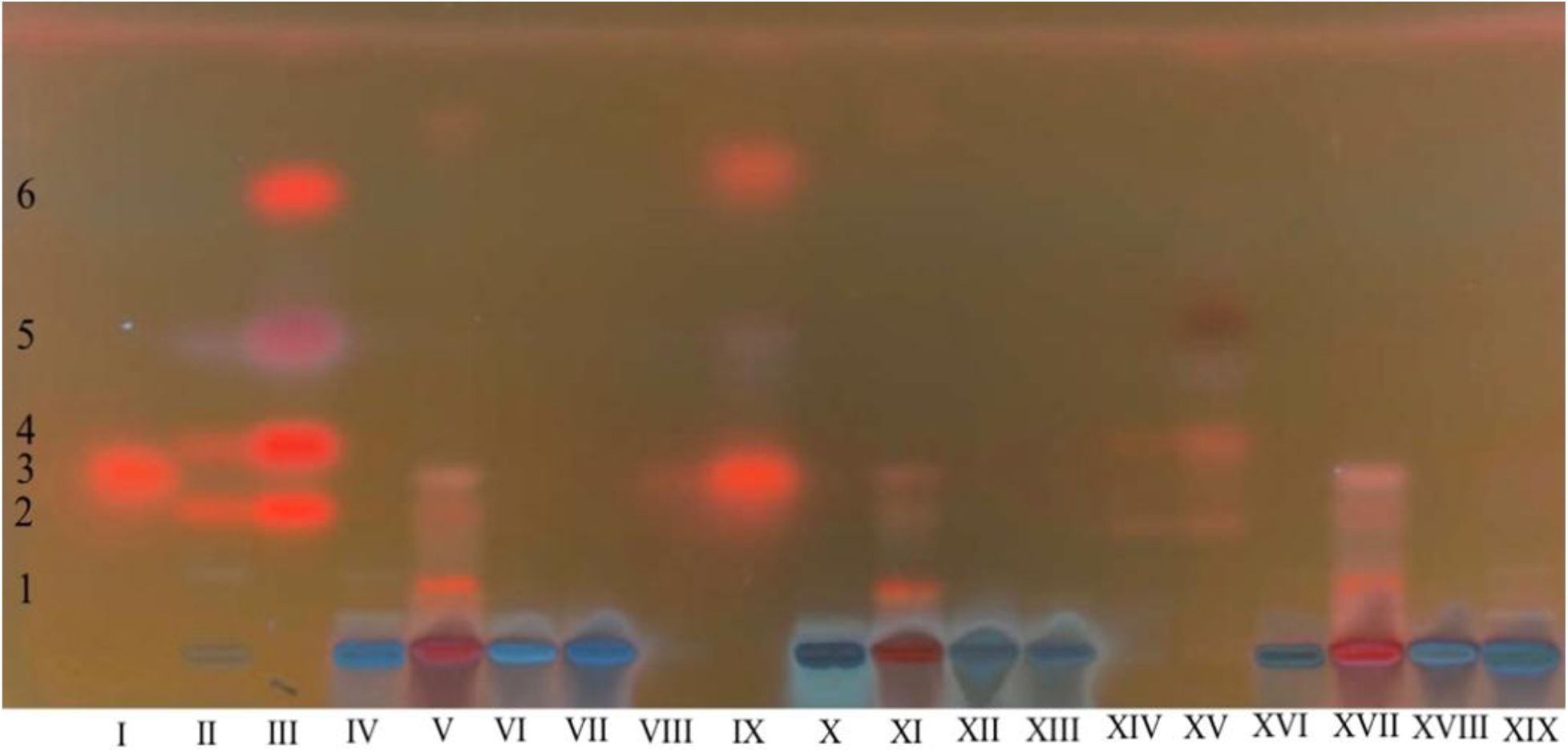
HPTLC analysis of mint samples. Arabic numerals represent bands, while roman numerals represent samples. I) Standard (Menthol) II) Simple Distillation of Spearmint III) Essence of Spearmint IV) Decoction of Simple Distillation of Spearmint V) Tincture of Spearmint VI) Infusion of Spearmint VII) Decoction of Spearmint VIII) Simple Distillation of Peppermint IX) Essence of Peppermint X) Decoction of Simple Distillation of Peppermint XI) Tincture of Peppermint XII) Infusion of Peppermint XIII) Decoction of Peppermint XIV) Simple Distillation of Unknown Mint XV) Essence of Unknown Mint XVI) Decoction of Simple Distillation of Unknown Mint XVII) Tincture of Unknown Mint XVIII) Infusion of Unknown Mint XIX) Decoction of Unknown Mint.

**Figure 3.**
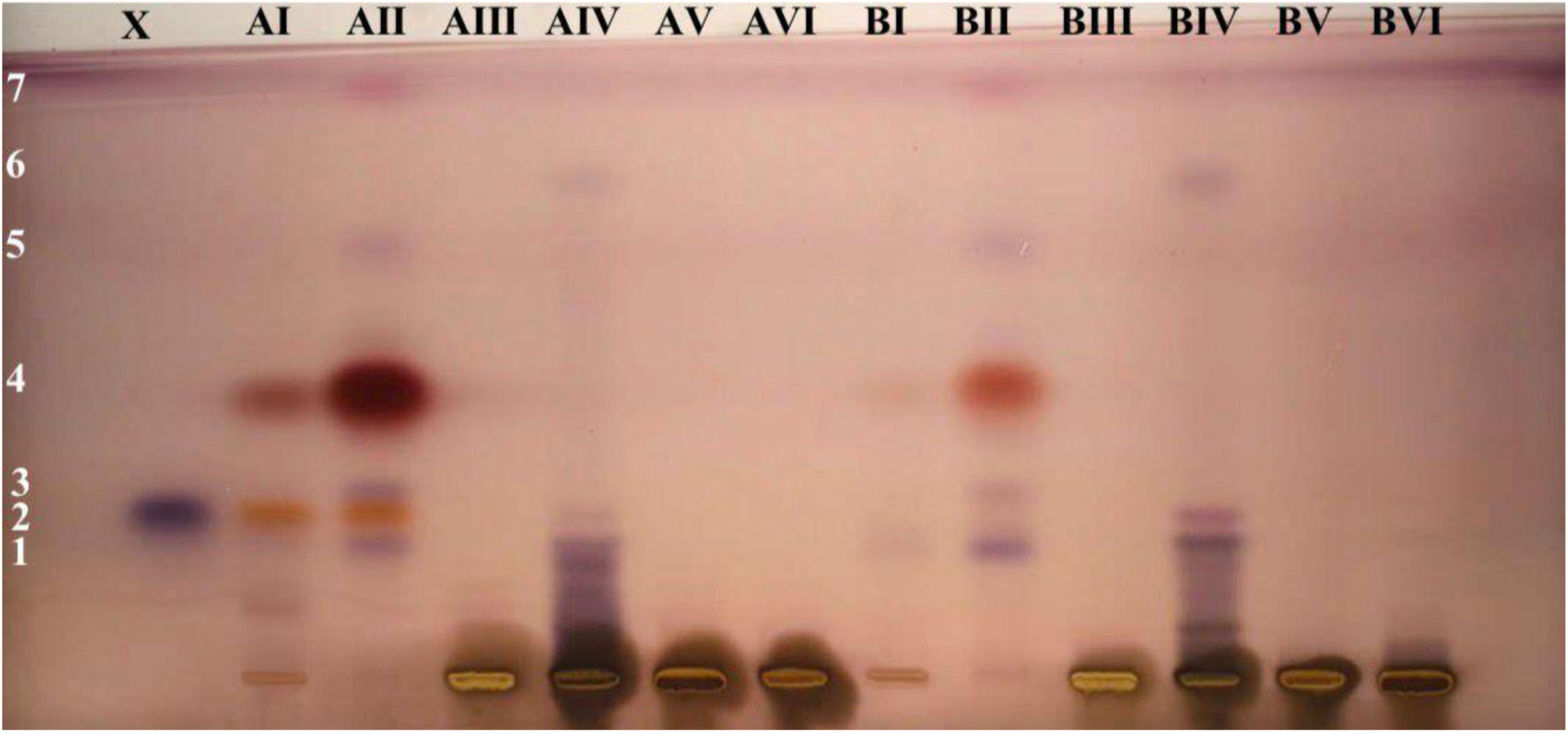
HPTLC of Mentha Longifolia from Bagh-e Firoozeh (A) and Shahr-e Rey (B). The Arabic numerals on the left represent the band numbers. The Roman numerals represent the different extraction methods as I) Simple Distillation II) Essence III) Decoction of Simple Distillation IV) Tincture V) Infusion VI) Decoction X) Standard (Menthol).

### 3.2 Simple Distillation

Simple distillation consistently yielded the least complex chromatographic profiles across all herbs. HPTLC plates showed either very few detectable bands or faint residues with low intensity (e.g., Figure 1, Lanes II, VIII, XIV; Figure 4, Lanes B, H, N). When present, bands corresponding to the thymol reference standard were weak and inconsistently observed. The limited number of bands suggests that simple distillation alone was inefficient in extracting a broad spectrum of phytochemicals, likely due to the loss of volatile compounds and insufficient solubilization of non-volatile constituents.

**Figure 4.**
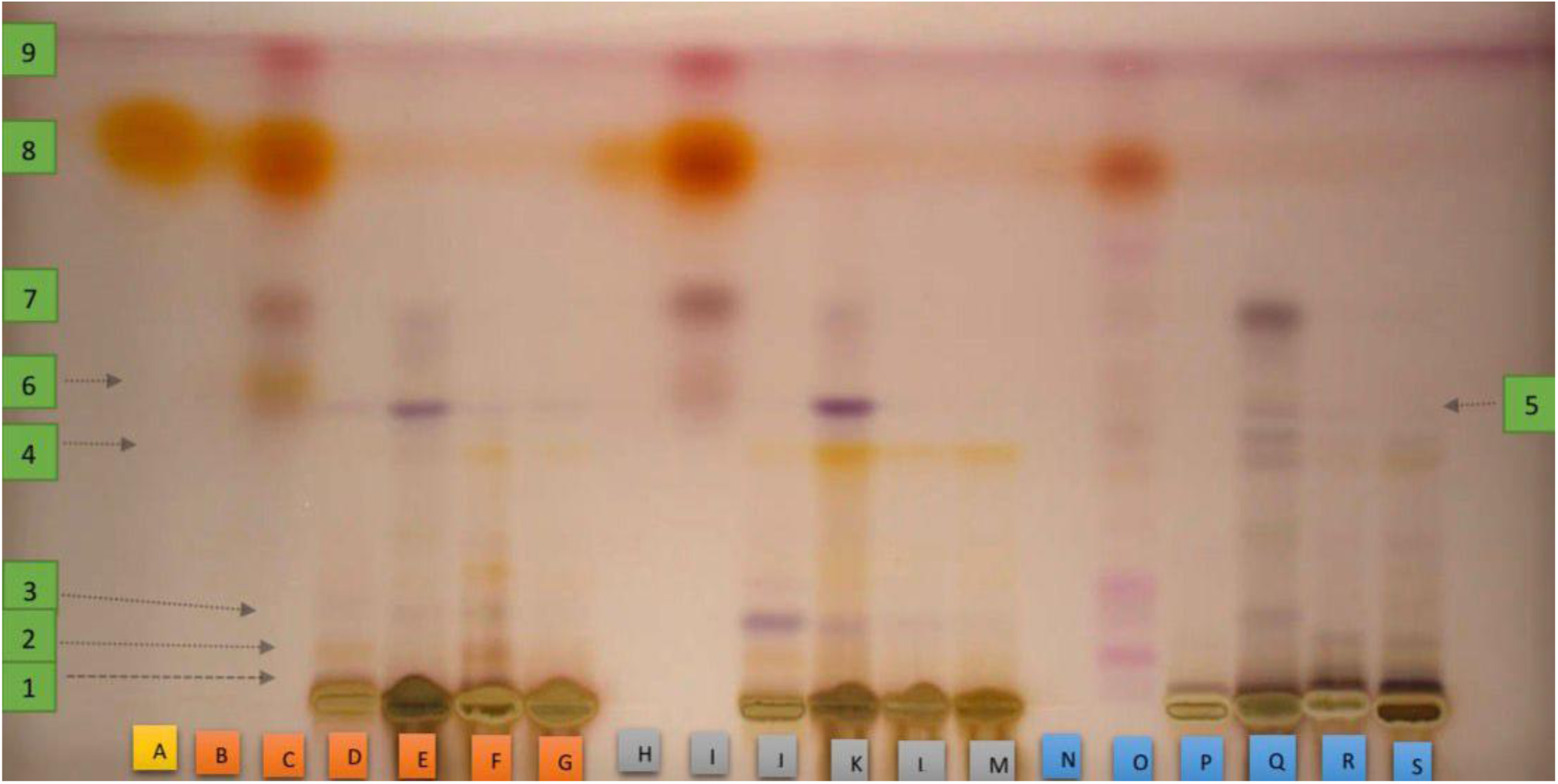
HPTLC Analysis of samples. Arabic numerals represent bands, while letters represent samples. A) Standard (Thymo). Letters B through G represent Thymus Vulgaris B) Simple Distillation C) Essence from simple distillation D) Decoction of simple distillation E) Tincture F) Infusion G) Decoction. Letters H through M represent Zataria Multiflora H) Simple Distillation I) Essence from simple distillation J) Decoction of simple distillation K) Tincture L) Infusion M) Decoction. Letters N through S represent Satureja Hortensis L N) Simple Distillation O) Essence from simple distillation P) Decoction of simple distillation Q) Tincture R) Infusion S) Decoction

### 3.3 Essential Oil (Essence) Extraction

Essential oil extracts exhibited the highest diversity of resolved bands among all extraction methods (e.g., Figure 1, Lanes III, IX, XV; Figure 4, Lanes C, I, O). These extracts consistently produced multiple high-Rf bands, indicating a predominance of non-polar and semi-polar compounds. Bands corresponding to thymol were clearly visible and well-resolved in most species, often appearing with high intensity (see Figure 4, Lane I). Although the exact band pattern varied among herbs, essential oil extracts consistently demonstrated broader phytochemical representation than aqueous methods. The concentration of bands in the upper regions of the HPTLC plates highlights the effectiveness of this method in isolating volatile terpenoids and lipophilic compounds. Among all extraction techniques, essential oil extraction produced the most reproducible non-polar profiles across different plant species.

### 3.4 Decoction of Distillate

The decoction of the residual aqueous phase obtained after simple distillation produced chromatographic profiles dominated by low-Rf bands. These bands were generally faint to moderately visible and were concentrated near the origin of the plate, indicating the presence of polar compounds (e.g., Figure 2, Lanes IV, X, XVI). Compared to simple distillation alone, decoction of distillate yielded a slightly higher number of detectable bands; however, overall diversity remained limited relative to essential oil and tincture extracts. This method demonstrated moderate effectiveness in recovering polar constituents that were not transferred into the distillate during distillation, though the resulting profiles remained less complex than those produced by other extraction approaches.

### 3.5 Tincture (Hydroalcoholic Extraction)

Hydroalcoholic tinctures consistently produced the widest polarity range of compounds among all extraction methods. HPTLC profiles from tincture extracts showed both low-Rf polar bands and high-Rf non-polar bands, resulting in broad and continuous band distributions across the plate (e.g., Figure 2, Lanes V, XI, XVII; Figure 4, Lanes E, K, Q). Band intensity in tincture extracts was generally higher than that observed in infusion and decoction, and comparable to essential oil extracts for certain compounds. While thymol-related bands were visible in several tincture samples, tinctures also revealed additional bands absent from essential oil extracts, indicating effective extraction of less volatile and moderately polar constituents. Overall, tincture extraction demonstrated superior versatility by capturing compounds spanning a wide polarity spectrum.

### 3.6 Infusion

Infusion extracts produced chromatographic profiles dominated by polar compounds. Bands were primarily located near the origin of the HPTLC plates, with low Rf values and limited upward migration (e.g., Figure 2, Lanes VI, XII, XVIII). Although some species exhibited a small number of additional bands with moderate Rf values, overall band diversity and intensity were reduced compared to essential oil and tincture extracts. The reproducibility of polar bands across different herbs suggests that infusion reliably extracts water-soluble constituents; however, the scarcity of higher-Rf bands indicates limited effectiveness in recovering volatile or lipophilic compounds.

### 3.7 Decoction

Decoction extracts yielded the simplest profiles among aqueous methods. HPTLC analysis revealed few detectable bands, most of which were highly polar and located close to the origin (e.g., Figure 4, Lanes G, M, S). In several cases, band intensity was weaker than that observed in infusion extracts, suggesting that prolonged heating may have contributed to thermal degradation or loss of certain phytochemicals. The reduced complexity of decoction profiles highlights its limited suitability for broad phytochemical screening when compared to other extraction techniques evaluated in this study.

### 3.8 Comparative Summary of Extraction Methods

When ranked according to qualitative phytochemical diversity and polarity coverage, extraction methods followed the general trend:

Essential oil ≈ Tincture > Infusion > Decoction of distillate > Decoction > Simple distillation

Essential oil and tincture extracts consistently outperformed aqueous and distillation-based methods in terms of band number, intensity, and polarity range. Aqueous extractions primarily yielded polar compounds with limited diversity, while simple distillation alone demonstrated minimal extraction efficiency.

A qualitative comparison of extraction methods is summarized in Table 1.

**Table 1.** Summary

| Extraction Method | Qualitative Diversity (Band Count) | Band Intensity | Dominant Rf Region / Compound Polarity | Key Findings & Extraction Mechanism |
| --- | --- | --- | --- | --- |
| Essential Oil (Essence) | Highest diversity | High | High-Rf (Non-polar and Semi-polar) | Most effective for isolating volatile and lipophilic compounds (e.g., Anethole, Menthol, Thymol). Most reproducible non-polar profiles. |
| **Tincture**<br>**(Hydroalcoholic)** | Highest diversity,<br>widest range | High, comparable<br>to Essential Oil for<br>some compounds | Broad spectrum<br>(Low-Rf and High-Rf) | Optimal for a<br>comprehensive<br>profile, effectively<br>extracting polar,<br>semi-polar, and<br>moderately non-polar compounds<br>simultaneously. |
| **Infusion** | Reduced diversity | Reduced<br>compared to<br>Essential<br>Oil/Tincture | Low-Rf (Polar) | Reliably extracts<br>water-soluble<br>constituents.<br>Limited recovery<br>of<br>volatile/lipophilic<br>compounds. |
| **Decoction of**<br>**Distillate** | Limited diversity | Faint to<br>moderately visible | Low-Rf (Polar) | Moderate<br>effectiveness in<br>recovering polar,<br>non-volatile<br>constituents that<br>remained in the<br>aqueous phase<br>after distillation. |
| **Decoction** | Simplest profiles<br>among aqueous<br>methods | Weaker than<br>Infusion | Low-Rf (Highly<br>Polar) | Least suitable for<br>broad screening.<br>Prolonged heating<br>may cause<br>thermal<br>degradation or loss of<br>phytochemicals. |
| **Simple**<br>**Distillation**<br>**(Aqueous**<br>**Distillate)** | Least complex<br>profiles | Very<br>low/inconsistent | Very Low-Rf<br>(Highly Polar) | Inefficient alone;<br>insufficient<br>solubilization of<br>non-volatile parts<br>and poor<br>retention of<br>volatiles in the<br>aqueous distillate. |

## 4. Discussion

### 4.1 Dominance of Extraction Method over Botanical Variation

The results of this study demonstrate that extraction method is the primary determinant of the qualitative phytochemical profiles observed by HPTLC, exceeding the influence of botanical differences among the investigated herbs. Although species-specific variation in band intensity and prominence was evident, the overall distribution of compounds, particularly in terms of polarity and diversity, was strongly governed by the extraction strategy employed. This finding supports the hypothesis that physicochemical extraction parameters such as solvent polarity, volatility, and thermal exposure exert a more pronounced effect on recovered phytochemicals than plant taxonomy when chemically compatible species are analyzed under standardized conditions. Aromatic herbs across the families *Lamiaceae*, *Apiaceae*, and *Asteraceae* share overlapping classes of secondary metabolites, including terpenoids and phenolic derivatives. This compositional overlap likely reduced interspecies variability and amplified extraction-dependent trends. Consequently, extraction method emerged as the dominant variable shaping HPTLC band patterns, consistent with previous observations that solvent and process selection critically influence phytochemical detectability (Sharma & Anand, 1997; Srivastava, 2011).

### 4.2 Mechanistic Interpretation of Method-Specific Profiles

The superior performance of essential oil extraction in generating diverse and intense high-Rf bands can be attributed to its effectiveness in isolating volatile and lipophilic compounds. Terpenoids such as thymol, menthol, and anethole exhibit low polarity and high vapor pressure, making them particularly amenable to distillation-based oil separation. The concentration of non-polar compounds in essential oil extracts explains their consistent migration to higher Rf regions and their clear visualization following anisaldehyde-sulfuric acid derivatization (Borugă et al., 2014; Meeran et al., 2017).

Hydroalcoholic tincture extraction produced the broadest polarity spectrum among all methods, reflecting the intermediate solvent polarity of ethanol-water mixtures. This solvent system facilitates the simultaneous extraction of polar, semi-polar, and moderately non-polar compounds, resulting in complex chromatographic profiles spanning a wide Rf range. The versatility of tincture extraction is particularly evident in its ability to recover compounds absent from both essential oil and aqueous extracts, underscoring its widespread use in pharmaceutical and herbal preparations (Polak et al., 2019).

In contrast, aqueous extraction methods such as, infusion and decoction, favored the recovery of polar constituents, as indicated by the predominance of low-Rf bands. The limited representation of non-polar compounds in these extracts reflects the low solubility of lipophilic metabolites in water. Additionally, prolonged heating during decoction may contribute to thermal degradation or volatilization of certain compounds, explaining the reduced band diversity observed relative to infusion. Simple distillation alone demonstrated minimal extraction efficiency, likely due to insufficient retention of volatile compounds in the aqueous distillate and poor solubilization of non-volatile constituents. The modest increase in polar compound recovery observed in decoctions of the distillate further supports the notion that distillation separates volatile compounds from the aqueous phase rather than concentrating them within a single extract.

### 4.3 Implications for Herbal Analysis and Applied Fields

The findings of this study have important implications for phytochemical screening, herbal medicine formulation, and industrial applications. For analytical purposes, extraction methods that yield broader and more representative phytochemical profiles such as essential oil extraction and hydroalcoholic tincture, are preferable for qualitative screening and comparative studies. These methods provide greater insight into the chemical complexity of plant materials and enhance detectability in chromatographic analyses. In pharmaceutical and cosmetic contexts, the polarity distribution of extracted compounds plays a crucial role in bioavailability, stability, and formulation performance. Essential oil extracts, rich in non-polar compounds, may be better suited for topical and aromatic applications, whereas tinctures offer greater flexibility by encompassing compounds with varied solubility and biological activity. Conversely, aqueous preparations may be appropriate when targeting polar constituents or when safety and simplicity are prioritized, albeit at the expense of chemical diversity.

### 4.4 Methodological Strengths and Limitations

A key strength of this study lies in its multi-species design and standardized analytical conditions, which allowed extraction-dependent trends to be evaluated independently of botanical variability. The use of HPTLC enabled direct visual comparison of multiple extracts on a single plate, enhancing reproducibility and comparative clarity. However, several limitations should be acknowledged. The qualitative nature of HPTLC analysis precludes precise quantification of individual compounds and limits conclusions regarding absolute extraction efficiency. Additionally, the use of a single mobile phase restricted optimization for compounds of varying polarity. Finally, the absence of compound identification techniques such as gas chromatography–mass spectrometry (GC-MS) prevented definitive assignment of individual bands.

### 4.5 Future Perspectives

Future studies could build upon these findings by incorporating quantitative densitometric analysis, multiple solvent systems, or hyphenated techniques such as HPTLC-MS or GC-MS to enable compound identification. Such approaches would complement the qualitative trends observed here and provide deeper insight into extraction efficiency and chemical composition.

## 5. Conclusion

This study demonstrates that extraction method is the principal factor governing the qualitative phytochemical profiles of aromatic medicinal herbs when assessed by high-performance thin-layer chromatography (HPTLC). Across ten herbs spanning three botanical families, chromatographic patterns clustered more consistently by extraction technique than by plant species, indicating that solvent polarity, volatility, and thermal conditions exert a stronger influence on observable phytochemistry than botanical variation under standardized analytical conditions.

Essential oil and hydroalcoholic tincture extractions consistently yielded the greatest diversity of resolved compounds and the widest polarity range, highlighting their suitability for comprehensive phytochemical screening. In contrast, aqueous methods such as infusion and decoction primarily recovered polar constituents and exhibited reduced band diversity, while simple distillation alone showed minimal extraction efficiency. These findings underscore the importance of method selection in phytochemical analysis and reinforce the role of extraction strategy as a critical determinant of analytical outcomes.

Overall, the results provide a comparative framework for selecting extraction techniques in herbal research, quality control, and applied fields such as pharmaceutical and cosmetic development. When qualitative chemical diversity and representative profiling are the primary objectives, extraction methods that accommodate a broad polarity spectrum are clearly favored.

## 6. Article Elements

### 6.1 Summary and Data

### 6.2 Author Contributions

AA focused on *Artemisia Dracunculus*, *Foeniculum Vulgare*, and *Anethum Graveolens L*. DF focused on *Mentha Piperita* and *Mentha Spicata*. AK focused on *Mentha Longifolia*. FG focused on *Satureja Hortensis L*, *Zataria Multiflora*, *Thymus Vulgaris*. All authors approved the final version.

### 6.3 Data Availability Statement

The raw chromatographic data (HPTLC plate images) and processed results are available from the corresponding author upon reasonable request.

## Acknowledgments

The authors would like to thank the staff and faculty of the Central and Herbal Medicine Laboratories at Shahid Beheshti University of Medical Sciences for providing access to the necessary equipment and facilities. This research received no external funding.

